# Spatiotemporal modulation of a fixed set of muscle synergies during unpredictable and predictable gait perturbations in older adults

**DOI:** 10.1101/2023.05.22.541808

**Authors:** Leon Brüll, Alessandro Santuz, Falk Mersmann, Sebastian Bohm, Michael Schwenk, Adamantios Arampatzis

## Abstract

Muscle synergies as functional low-dimensional building blocks of the neuromotor system regulate the activation patterns of muscle groups in a modular structure during locomotion. The purpose of the current study was to explore how older adults organize locomotor muscle synergies to counteract unpredictable and predictable gait perturbations during the perturbed and the recovery steps. Sixty-three healthy older adults (71.2 ± 5.2 years) participated in the study. Mediolateral and anteroposterior unpredictable and predictable perturbations during walking were introduced using a treadmill. Muscle synergies were extracted from the electromyographic activity of 13 lower limb muscles using Gaussian non-negative matrix factorization. The four basic synergies responsible for unperturbed walking (weight acceptance, propulsion, early swing and late swing) were preserved in all applied gait perturbations, yet their temporal recruitment and muscle contribution in each synergy were modified (p<0.05). These modifications were observed up to four recovery steps and were more pronounced (p<0.05) following unpredictable perturbations. The recruitment of the four basic walking synergies in the perturbed and recovery gait cycles indicates a robust neuromotor control of locomotion by using activation patterns of a few and well-known muscle synergies with specific adjustments within the synergies. The selection of pre-existing muscle synergies while adjusting the time of their recruitment during challenging locomotor conditions may facilitate the effectiveness to deal with perturbations and promote the transfer of adaptation between different kinds of perturbations.

**Summary statement:** The flexible recruitment of the four and well-known muscle synergies responsible for unperturbed walking during unpredictable and predictable gait perturbations indicates an effective way to counteract locomotor perturbations, where fast reactive responses are necessary to maintain postural stability.

## Introduction

Investigating neuromotor control during gait perturbations in older adults might help to better understand real-life mobility in the elderly. After an unpredictable gait perturbation, more than five recovery steps can be required to return to baseline walking stability (Epro et al., 2018; Oddsson et al., 2004). Fast and effective motor responses in order to maintain stability are of high importance after an unpredictable perturbation (Karamanidis et al., 2020; Maki & McIlroy, 2006), since beneficial mechanisms modulated by anticipation and prediction (Patla, 2003) are not available. Short latency reflexes, 45 – 70 ms, possibly mediated by subcortical centers (McDonagh & Duncan, 2002; Quintern et al., 1985; Van Der Linden et al., 2007), rapidly generate neuromuscular responses after a locomotor perturbation, triggering postural corrections (Cronin et al., 2009; Nieuwenhuijzen & Duysens, 2007). However, the short monosynaptic spinal feedback loops such as the stretch and the Golgi tendon reflexes are relatively unaffected by prior experience (Nieuwenhuijzen & Duysens, 2007; Rothwell et al., 1986). On the other hand, longer feedback loops such as long latency reflexes which involve supraspinal mechanisms (Taube et al., 2006) can be modified by prior experience to enhance the effectiveness of the task-specific neuromuscular output (Wolpert & Landy, 2012). Experience-based learning leads to sensorimotor adaptations, which are supposedly mediated by the cerebellum (Diedrichsen et al., 2005; Morton & Bastian, 2006). Prior experience with a certain type of perturbation enhances the opportunity to proactively plan the desired movement with ongoing adaptative adjustments in order to reduce the consequences of the perturbation (Bierbaum et al., 2010; Marigold & Patla, 2002; Pai et al., 2003).

Muscle synergies (Bizzi et al., 1991) have been frequently used to investigate the neuromotor control of human locomotion in unperturbed and perturbed conditions (Cappellini et al., 2006; Dominici et al., 2011; Santuz et al., 2018a; Ting et al., 2015). Muscle synergies as functional low-dimensional building blocks of the neuromotor system regulate the activation patterns of muscle groups and coordinate the complex behavior of human locomotion in a modular structure (Dominici et al., 2011; Ivanenko et al., 2004). There is a linkage between muscle synergies and the connectivity of spinal premotor interneurons as well as motor cortical neurons (Amundsen Huffmaster et al., 2018; Hart & Giszter, 2010; Takei et al., 2017), which provides evidence that muscle synergies are important neurophysiological entities for motor control and learning (Bizzi & Cheung, 2013). Muscle synergies as neural control modules may be shared across a large repertoire of motor tasks (Chvatal & Ting, 2013; D’Avella & Bizzi, 2005) and flexibly recruited by descending neural pathways from supraspinal regions (Roh et al., 2011), which would be an efficient strategy to generate complex motor behavior. The modular architecture of muscle synergies simplifies motor control and facilitates an effective execution of multiple motor tasks (D’Avella et al., 2003; D’Avella & Bizzi, 2005). It has been found that the spatiotemporal structure and composition of a small number of muscle synergies is generated robustly to account for the changing biomechanical demands of the task and alterations of the neural and musculoskeletal systems during locomotion (Cheung et al., 2020; Yang et al., 2019).

Four basic locomotor muscle synergies associated with distinct gait phases (i.e., weight acceptance, propulsion, early swing and late swing) were reported for steady-state unperturbed walking (Clark et al., 2010; Dominici et al., 2011; Lacquaniti et al., 2012; Santuz et al., 2018a). In the presence of consecutive external mechanical perturbations, as for example during walking on slippery or uneven terrains, the structure and number of the muscle synergies is retained, while a widening and variable recruitment of their basic activation patterns serves as a compensatory strategy to deal with the perturbations (Martino et al., 2015; Santuz et al., 2018a). Chvatal and Ting (2012) found that a fixed set of muscle synergies could explain the muscle activation patterns of the perturbed step during both anteroposterior and mediolateral gait perturbations. On the other hand, Oliveira et al. (2012) and Nazifi et al. (2017) reported that unperturbed and perturbed walking have only a fraction of shared synergies and that the muscle weights of unperturbed walking only marginally agreed with the perturbed ones. All the above-mentioned studies investigated young participants with prior experience with the induced gait perturbations. Little is known on the spatiotemporal structure and composition of the muscle synergies and the influence of prior experience in older adults. Gait stability after unpredictable and novel perturbations is deteriorated by a greater degree in older adults than in young ones (Bierbaum et al., 2010; Maki & McIlroy, 2006). Furthermore, older adults usually need more recovery steps until complete recovery (Süptitz et al., 2013). Nevertheless, the potential to adapt based on prior experience is still maintained in older adults (Bierbaum et al., 2011; Bohm et al., 2015; Marigold & Patla, 2002).

A fast recruitment of muscle synergies related to specific functional phases of walking after a gait perturbation might be crucial for a successful recovery. The activation of a fixed set of well-known locomotor muscle synergies while adjusting the time of their recruitment may provide a more effective way to counteract locomotor perturbations than the generation of new muscle synergies. The purpose of the current study was to explore how older adults organize locomotor muscle synergies to counteract predictable and unpredictable gait perturbations during the perturbed and the recovery steps. For this purpose, we investigated their responses to anteroposterior and mediolateral perturbations with and without prior experience during walking on a treadmill and we measured the electromyographic (EMG) activity of thirteen muscles of the perturbed leg. We hypothesized that older adults would use the four basic muscle synergies of unperturbed walking (weight acceptance, propulsion, early swing and late swing) and adjust the time of their recruitment during the perturbed and the recovery steps. We further hypothesized that the spatiotemporal adjustments in muscle synergies would be less pronounced during predictable perturbations with prior experience compared to the unpredictable perturbations without prior experience.

## Methods

### Participants

Sixty-three healthy older adults (33 females, age 71.2 ± 5.2 years, height 171.3 ± 8.9 cm, body mass 71.6 ± 11.9 kg, means ± standard deviation) participated in this study. Inclusion criteria were an age of 65-80 years, home-dwelling and the ability to walk independently on a treadmill for at least 20 min. Participants were excluded if they had neurological, musculoskeletal or cardiovascular disorders, uncorrected visual impairments, severe vertigo or a body mass index ≥ 30 due to potential issues with the EMG recordings (Nordander et al., 2003). All participants gave their written consent to the experimental procedure. This study was reviewed and approved by the Ethics Committee of Heidelberg University (AZ Schw 2018 1/2).

### Experimental protocol

In the current study, all experiments were conducted during walking on a particular perturbation treadmill (BalanceTutor™, MediTouch LTD, Netanya, Israel, figure 1). The treadmill can introduce two types of perturbations (i.e. anteroposterior and mediolateral direction) during human gait. Mediolateral perturbations always consisted of a fast shift of 11.7 cm of the treadmill platform to the left (acceleration: 1.3 m/s², duration of acceleration: 0.3 s, duration of deceleration: 0.3 s). Anteroposterior perturbations consisted of an alteration of the speed of the belt of 0.82 m/s (acceleration: 4.1 m/s², duration of acceleration: 0.2 s, duration of deceleration: 0.2 s). For reasons of safety, participants were strapped to a harness system, which was set to prevent the knees from touching the ground in case of a fall. The participants were randomly divided into two groups (A and B), where each group experienced the perturbations in a different order with regard to the directions.

**Figure 1.**
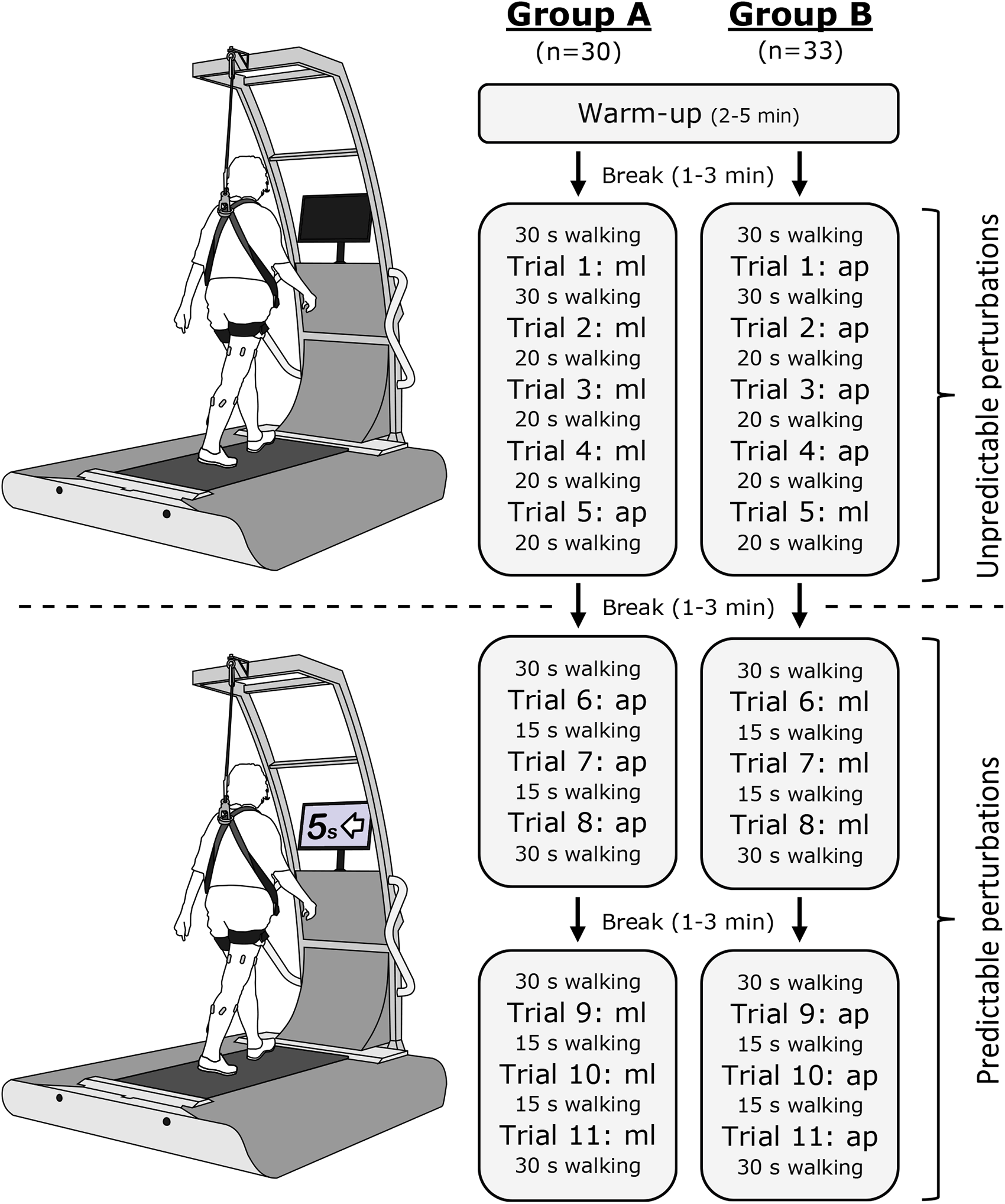
*Study design on the perturbation treadmill showing the protocols for both groups (A and B).* The protocols included a warm-up with unperturbed walking, a block of unpredictable perturbations and two blocks of predictable perturbations (screen showing a countdown and the direction of the upcoming perturbation) in the different directions. Trials 1, 5, 8 and 11 were selected for analysis. ml = mediolateral (i.e. a shift of the treadmill platform to the left), ap = anteroposterior (i.e. an acceleration of the treadmill belt).

After a familiarization session, which included two sets of 10 min unperturbed treadmill walking, the participants were invited to the main experiment on a separate day. The experiment started with a warm-up of 2 to 5 min walking with a gradual speed progression until the target speed of 1.1 m/s was reached. This speed was chosen based on normative data of the included age group (Hollman et al., 2011). In the case of an induced perturbation, the participants were requested to regain balance without stopping to walk. In the first perturbation protocol, we introduced an unpredictable perturbation (mediolateral for group A and anteroposterior for group B), which was unknown in time and type (figure 1). We continued the protocol with three perturbations of the same type that were still unannounced. Subsequently, an unpredictable perturbation for each group in the other direction appeared (anteroposterior for group A and mediolateral for group B). We finalized the protocol with three predictable anteroposterior and three predictable mediolateral perturbations for group A, and accordingly three predictable mediolateral and three predictable anteroposterior perturbations for group B. In the predictable perturbations, the time and direction of the perturbation was announced on a screen in front of the participants. In the analysis, we selected the first unknown perturbation per direction for the unpredictable condition (trials 1 and 5) and the third announced perturbation per direction (trials 8 and 11) for the predictable condition as the participants had already experience with this kind of perturbation. All perturbations were induced in mid-stance of the right leg, as detected by automatic elaboration of the data of four built-in force sensors (sampling frequency: 60 Hz) integrated in the perturbation treadmill.

For each of the four perturbed conditions (unpredictable mediolateral, predictable mediolateral, unpredictable anteroposterior, predictable anteroposterior) 12 consecutive gait cycles surrounding the selected perturbations were included into the analysis. This series of cycles included 3 cycles before the perturbation, the perturbation cycle and 8 cycles after the perturbed cycle. A preliminary analysis of the gait parameters (cadence, stance duration, swing duration and duty factor, defined as the fraction of the gait cycle time that the foot spends on the ground) revealed that up to 7 cycles were significantly affected by the perturbation.

### Measurements and data analysis

We recorded the EMG activity of 13 ipsilateral (right side) muscles of the lower limb at 2 kHz using a 16-channel wireless bipolar surface EMG system (aktos, myon AG, Schwarzenberg, Switzerland). The following muscles were included: gluteus medius, gluteus maximus, tensor fasciæ latæ, rectus femoris, vastus medialis, vastus lateralis, semitendinosus, biceps femoris, tibialis anterior, peroneus longus, gastrocnemius medialis, gastrocnemius lateralis and soleus. Foam-hydrogel electrodes with snap connector (H124SG, Medtronic plc, Dublin, Ireland) were attached according to SENIAM (Surface EMG for non-invasive assessment of muscles) recommendations (Hermens et al., 2000).

In order to detect the stance and swing phase of gait, we used a 3d-accelerometer (143 Hz) synchronized with the EMG system attached to the right shoe over the most proximal portion of the second to fourth metatarsal bones. The raw accelerometer data were filtered by a 4^th^-order infinite impulse response (IIR) Butterworth zero-phase low-pass filter with a 15 Hz cut-off frequency. To estimate touchdown, the modified version of the foot contact algorithm developed by Maiwald and colleagues was used (Maiwald et al., 2009). Briefly, the minimal vertical acceleration was found for each cycle and an interval of – 150 ms to +100 ms was built around it. Touchdown was then defined by a characteristic maximum in the vertical acceleration within that interval. For the estimation of lift-off, we developed a new algorithm, in which an interval was built from 300 ms after the touchdown event until 200 ms before the subsequent touchdown. Lift-off was then defined as the minimum of the anteroposterior acceleration of the foot within that interval. This approach was validated against data recorded with a force plate (900 x 600 mm, AMTI BP600, Advanced Mechanical Technology, Inc., Watertown, MA, USA) in a different sample (n=31). The average deviation in lift-off detection was 3 ± 12 ms. The average deviation of 3 ms was added to all detected lift-off events to achieve the highest possible accuracy.

From the EMG data, we extracted the muscle synergies for the investigated trials using a custom R script (R Core Team, 2021; R Foundation for Statistical Computing, Vienna, Austria) that utilizes the classical Gaussian non-negative matrix factorization (NMF) algorithm (Lee & Seung, 1999; Santuz et al., 2017). The raw EMG signals were filtered in the acquisition device by a band-pass filter with cut-off frequencies of 10 and 500 Hz. Then the signals were first high-pass filtered using a 4^th^-order IIR Butterworth zero-phase filter with a cut-off frequency of 50 Hz. Thereafter, the signals were full-wave rectified before a 4^th^-order IIR Butterworth zero-phase low-pass filter with a cut-off frequency of 20 Hz was applied. The amplitude of each single EMG channel (representing a specific muscle) was normalized to its maximum, after subtracting the minimum, in every trial. Each gait cycle of each condition (12 cycles x 4 conditions) was analyzed separately for each participant. To increase the robustness of the factorization, the NMF was repeated 20 times for each cycle and then averaged. Each gait cycle was then normalized in time to 200 points, where stance and swing phase each got an equal distribution of 100 points. This transformation was conducted to increase comprehension and interpretation due to improved comparability between different participants, conditions and gait phases (Santuz et al., 2018b).

The extraction of the muscle synergies through the NMF was conducted as previously reported by our group (Santuz et al., 2020; Santuz et al., 2018a). At first, a matrix *V = m* x *n* was created from the original data, where *m* is the 13 recorded muscle activities and *n* is the number of normalized time points (i.e. 200 x 12 cycles). This matrix was factorized utilizing the NMF so that *V* ≈ *VR* = *MP^T^*, where *VR* is a new approximation of the matrix *V*, which is reconstructed through a multiplication of the two matrices *M* and *P*. *P* contains the time-dependent activation patterns of a synergy (Dominici et al., 2011; Santuz et al., 2017) and have the dimensions *p × n*, where the number of rows *p* represents the minimum number of synergies necessary to satisfactorily reconstruct the original set of signals *V.* The matrix *M* contains the muscle weights, which are the time-invariant weights of the synergies (Gizzi et al., 2011; Santuz et al., 2017) and has the dimensions *m* × *p*. The quality of the reconstruction of the original data matrix through the extracted synergies was assessed utilizing the coefficient of determination R² as described earlier by Santuz et al. (2017).

Fundamental synergies, defined as activation patterns with a single main activation peak (Santuz et al., 2018a), were classified as described in detail by Munoz-Martel et al. (2021). In short, muscle synergies were classified using a method based on k-means clustering. Thirteen iterations (number of included muscles) were computed building one to 13 clusters from the activation patterns of all participants for every single cycle of each condition separately. For each iteration, the sum of squares within the contained clusters was determined. Then, a curve of the sum of squares versus the number of clusters was built. The minimum number of clusters before the curve fits a linear interpolation with a mean squared error below 10^-3^, was used as a criterion to determine the classification. Muscle synergies that were not classified (18.7% in the current study) were excluded from further analysis.

The cosine similarity (CS) of the muscle weights and activation patterns of the two cycles before the perturbation, the perturbed cycle and the following recovery cycles with regard to the first unperturbed cycle (i.e. third cycle before the perturbation) was calculated in order to assess the modifications in the spatiotemporal structure (muscle weights and activation patterns) of the muscles synergies during the experiment (formula 1). We used the third cycle before the perturbation as a reference to calculate the CS because it was the most remote from the perturbed cycle and therefore is defined as unperturbed walking.

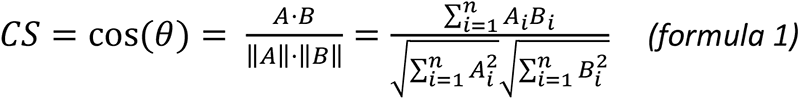

*A* represents the vector of the muscle weights (*n*=13) or activation patterns (*n*=200) of the third cycle before the perturbation and *B* the vector of the muscle weights or activation patterns of all following gait cycles. The values of CS range from 1 meaning exactly the same pattern to −1 meaning exactly the opposite.

To compare the activation patterns of the muscle synergies between conditions, we additionally used the full width at half maximum (FWHM) and the center of activity (CoA), which are established metrics for the width and timing of the basic activation patterns (Cappellini et al., 2016; Martino et al., 2014; Santuz et al., 2020). The FWHM was calculated for each cycle accumulating all points exceeding the half maximum of the activation patterns, after subtracting the minimum (Martino et al., 2014). The CoA was also calculated cycle-by-cycle as the angle of the vector in polar coordinates that points to the center of mass of that circular distribution (Cappellini et al., 2016). The polar direction represented the cycle’s phase, with angle 0 ≤ θ ≤ 2π.

### Statistics

For the parameters cadence, stance time, swing time, duty factor, FWHM, CoA and CS a linear mixed-effects model was used with predictability (unpredictable, predictable) and gait cycle as fixed factors and participant as a random factor. The modeling was conducted using the function “lme” of the R package “nlme” v 3.1-152 (Pinheiro & Bates, 2021), with maximizing the restricted log-likelihood and for both perturbation directions (mediolateral or anteroposterior) separately. In case of significant effects of a factor or an interaction between factors predictability and gait cycle, a *post-hoc* analysis was performed comparing all included cycles with the third cycle prior to the perturbation as the one with the higher temporal distance to the upcoming perturbation. For the CS, the *post-hoc* comparisons were performed with regard to the CS of the first and second unperturbed cycles as a representative metric for unperturbed walking. For all analyzed parameters, differences between unpredictable and predictable cycles were investigated by pairwise comparisons of the matching cycles (e.g. cycle 1 in unpredictable vs. cycle 1 in predictable). The p-values for all *post-hoc* tests were adjusted for multiple comparisons according to Benjamini-Hochberg. All these analyses were conducted in R v4.1.0.

To check for a proactive modulation of muscle activity in the predictable trials, statistical parametric mapping (SPM) paired two-tailed t-tests (Pataky, 2010) were performed on the muscle’s EMG-activity during the stance phase comparing the predictable and unpredictable conditions for both mediolateral and anteroposterior perturbations. The initiation of the perturbation occurred in the middle of the stance phase, thus any proactive adjustments of muscle activity in the predictable trials might be visible in the EMG-activity during the first half of the stance phase. For the SPM, the EMG-activity of each muscle was normalized to the maximum activity during unperturbed walking (specifically to the third cycle prior to the perturbation). The SPM was conducted in MATLAB R2017b (The MathWorks Inc., MA, United States) using the spm1d-package (version M.0.4.10). The level of significance was set at α = 0.05 for all analyses.

## Results

We found a significant effect of predictability (p < 0.001) and gait cycle (p < 0.001) on cadence, stance time, swing time and duty factor in both mediolateral and anteroposterior perturbations (table 1). The post-hoc analysis showed that all four temporal gait parameters were significantly affected by the perturbation in at least one and up to seven cycles (p < 0.05) in each of the four perturbed conditions compared to the respective unperturbed gait cycle (table 1). The three cycles before the perturbation did not show any statistically significant differences (p = 0.228 to 0.998) in the temporal parameters in either condition (table 1). Cadence generally increased, while stance and swing durations decreased due to the perturbation. The duty factor increased in the perturbed cycle but decreased in the following recovery cycles compared to unperturbed walking. The observed alterations in the gait parameters during the unpredictable condition were significantly higher in both perturbation directions in two and up to seven recovery gait cycles (p < 0.05) compared to the predictable one (table 1).

**Table 1.** Means ± SD values of cadence, stance time, swing time and duty factor during the three unperturbed cycles (U1 to U3), the perturbation cycle (P) and the following 8 recovery cycles (R1 to R8) for the four different conditions. Statistically significant differences to the first unperturbed cycle (*) and between the unpredictable and predictable condition (†) are highlighted (p<0.05).

|  |  |  | U1 | U2 | U3 | P | R1 | R2 | R3 | R4 | R5 | R6 | R7 | R8 |
| --- | --- | --- | --- | --- | --- | --- | --- | --- | --- | --- | --- | --- | --- | --- |
| Mediolateral perturbation | Unpredictable | Cadence [steps/min] | 109.9 ± 7.0 | 110.0 ± 7.0 | 109.9 ± 6.3 | 128.6 ± 10.7**† | 161.8 ± 36.3**† | 134.1 ± 22.2**† | 124.1 ± 17.1*† | 118.2 ± 13.7*† | 115.3 ± 10.7**† | 113.6 ± 9.0 | 113.0 ± 8.8 | 112.0 ± 7.8 |
|  |  | Stance time [ms] | 702 ± 53 | 702 ± 54 | 704 ± 48 | 685 ± 55 | 454 ± 104**† | 541 ± 105**† | 608 ± 94**† | 643 ± 79**† | 661 ± 66**† | 676 ± 62**† | 681 ± 58**† | 689 ± 57 |
|  |  | Swing time [ms] | 395 ± 25 | 393 ± 25 | 392 ± 27 | 254 ± 44**† | 321 ± 71**† | 377 ± 47**† | 375 ± 39**† | 384 ± 36 | 388 ± 32 | 387 ± 30 | 386 ± 29 | 387 ± 29 |
|  |  | Duty factor | 0.64 ± 0.02 | 0.64 ± 0.02 | 0.64 ± 0.02 | 0.73 ± 0.03**† | 0.59 ± 0.05**† | 0.59 ± 0.04**† | 0.62 ± 0.03**† | 0.63 ± 0.03**† | 0.63 ± 0.02**† | 0.64 ± 0.02 | 0.64 ± 0.02 | 0.64 ± 0.02 |
|  | Predictable | Cadence [steps/min] | 110.2 ± 6.7 | 110.7 ± 7.2 | 111.3 ± 7.6 | 121.6 ± 12.8**† | 119.2 ± 14.7**† | 112.8 ± 9.3† | 111.4 ± 7.1† | 110.9 ± 6.9† | 110.8 ± 6.6† | 110.5 ± 6.7 | 110.9 ± 6.8 | 110.3 ± 6.5 |
|  |  | Stance time [ms] | 699 ± 51 | 694 ± 53 | 689 ± 57 | 670 ± 55* | 632 ± 87**† | 672 ± 62**† | 688 ± 54† | 692 ± 56† | 697 ± 51† | 698 ± 51† | 699 ± 53† | 0.70 ± 0.05 |
|  |  | Swing time [ms] | 393 ± 31 | 395 ± 32 | 394 ± 33 | 328 ± 64**† | 389 ± 41† | 398 ± 35† | 394 ± 33† | 394 ± 30 | 389 ± 32 | 394 ± 32 | 389 ± 30 | 393 ± 29 |
|  |  | Duty factor | 0.64 ± 0.02 | 0.64 ± 0.02 | 0.64 ± 0.02 | 0.67 ± 0.04**† | 0.62 ± 0.03**† | 0.63 ± 0.02**† | 0.64 ± 0.02† | 0.64 ± 0.02† | 0.64 ± 0.02† | 0.64 ± 0.02 | 0.64 ± 0.02 | 0.64 ± 0.02 |
| Anteroposterior perturbation | Unpredictable | Cadence [steps/min] | 109.4 ± 6.8 | 109.2 ± 6.5 | 109.5 ± 6.5 | 130.2 ± 8.6* | 150.3 ± 36.3**† | 121.4 ± 15.2**† | 115.7 ± 13.0**† | 114.5 ± 12.3* | 112.9 ± 11.6 | 112.2 ± 10.7 | 111.9 ± 11.4 | 112.1 ± 12.1 |
|  |  | Stance time [ms] | 706 ± 52 | 705 ± 51 | 705 ± 51 | 627 ± 49* | 515 ± 136**† | 617 ± 89**† | 663 ± 74**† | 668 ± 70**† | 680 ± 64* | 688 ± 62 | 690 ± 64 | 689 ± 70 |
|  |  | Swing time [ms] | 396 ± 28 | 398 ± 30 | 394 ± 26 | 299 ± 38**† | 324 ± 55**† | 386 ± 34* | 385 ± 32* | 389 ± 31 | 391 ± 30 | 389 ± 30 | 391 ± 31 | 390 ± 30 |
|  |  | Duty factor | 0.64 ± 0.02 | 0.64 ± 0.02 | 0.64 ± 0.02 | 0.68 ± 0.03**† | 0.61 ± 0.05* | 0.61 ± 0.03**† | 0.63 ± 0.03 | 0.63 ± 0.02† | 0.63 ± 0.02 | 0.64 ± 0.02 | 0.64 ± 0.02 | 0.64 ± 0.02 |
|  | Predictable | Cadence [steps/min] | 110.7 ± 7.6 | 110.6 ± 7.6 | 111.5 ± 8.4 | 129.7 ± 11.1* | 121.5 ± 20.6**† | 111.9 ± 9.5† | 111.6 ± 7.5† | 111.0 ± 7.7 | 111.1 ± 7.7 | 110.7 ± 7.8 | 110.7 ± 7.4 | 110.2 ± 7.2 |
|  |  | Stance time [ms] | 696 ± 55 | 697 ± 55 | 690 ± 61 | 611 ± 55* | 625 ± 117**† | 691 ± 69† | 687 ± 58† | 697 ± 59† | 694 ± 55 | 695 ± 56 | 697 ± 54 | 701 ± 54 |
|  |  | Swing time [ms] | 393 ± 29 | 392 ± 28 | 391 ± 28 | 320 ± 32**† | 386 ± 41† | 389 ± 27 | 393 ± 28 | 389 ± 28 | 391 ± 29 | 394 ± 30 | 392 ± 28 | 393 ± 30 |
|  |  | Duty factor | 0.64 ± 0.02 | 0.64 ± 0.02 | 0.64 ± 0.02 | 0.66 ± 0.02**† | 0.61 ± 0.04* | 0.64 ± 0.02† | 0.64 ± 0.02 | 0.64 ± 0.02† | 0.64 ± 0.02 | 0.64 ± 0.02 | 0.64 ± 0.02 | 0.64 ± 0.02 |

An example of the EMG-activity in the unperturbed, perturbed and recovery gait cycles of every recorded muscle is presented in figure 2. In all gait cycles of all conditions, we identified four muscle synergies that were able to sufficiently reconstruct the recorded EMG data (R² was 0.89 ± 0.03 for mediolateral unpredictable, 0.88 ± 0.03 for mediolateral predictable, 0.90 ± 0.03 for anteroposterior unpredictable and 0.89 ± 0.02 for anteroposterior predictable). The four synergies are functionally associated with the phases of the gait, namely the weight acceptance synergy with high activation of knee extensors and glutei, the propulsion synergy with a main contribution of plantar flexors, the early swing synergy mainly represented by the foot dorsiflexors and the late swing synergy highly related to the knee flexors (figure 3).

**Figure 2.**
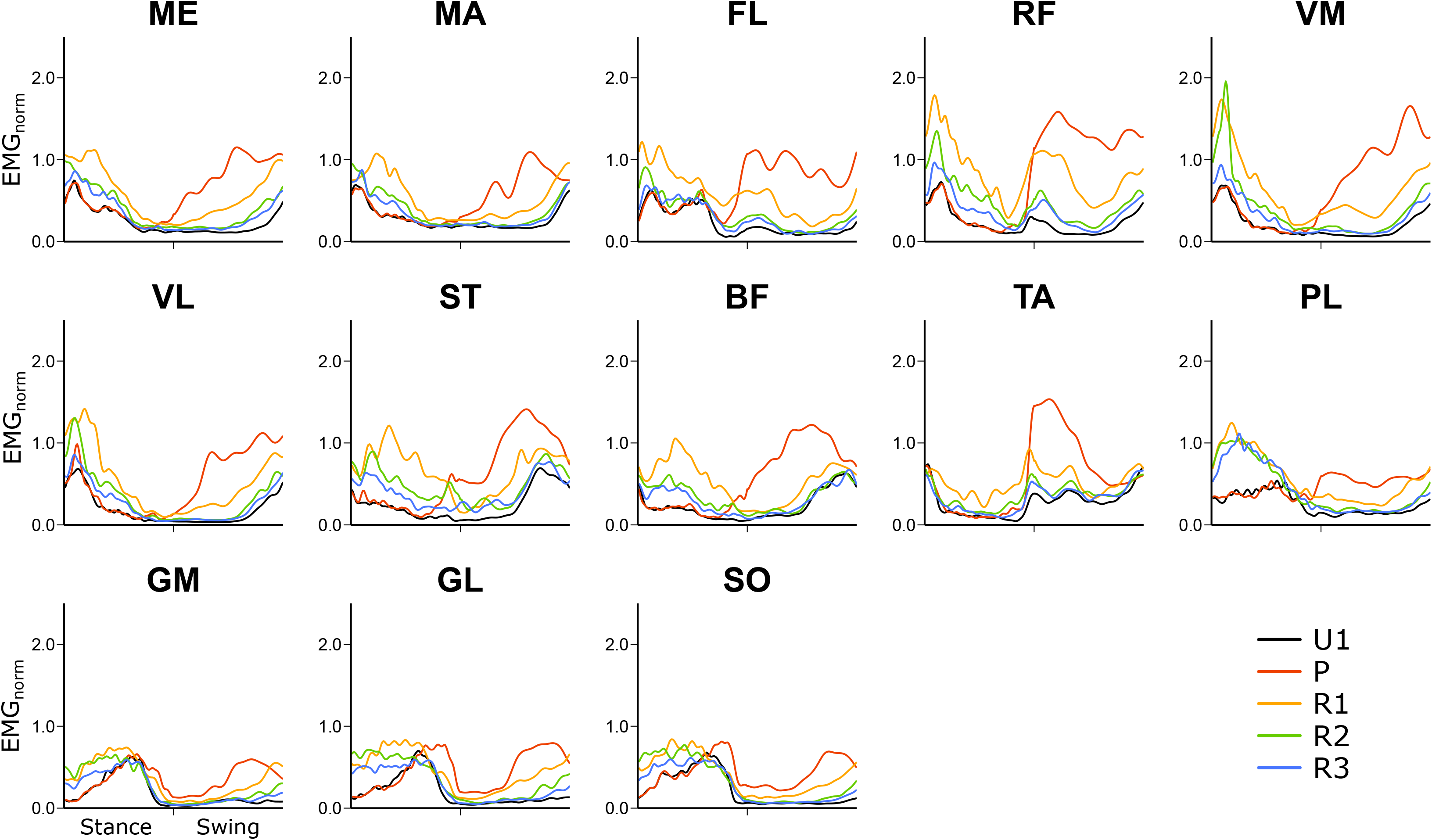
Means curves of the electromyographic (EMG) activity of the 13 recorded leg muscles in the mediolateral unpredictable condition during the unperturbed walking (U1), the perturbation cycle (P) and the following three recovery cycles (R1 to R3). For each muscle, the amplitude was normalized to the maximum EMG activity during unperturbed walking. The stance and swing phases were normalized to an equal distribution of data points. ME = gluteus medius, MA = gluteus maximus, FL = tensor fasciæ latæ, RF = rectus femoris, VM = vastus medialis, VL = vastus lateralis, ST = semitendinosus, BF = biceps femoris, TA = tibialis anterior, PL = peroneus longus, GM = gastrocnemius medialis, GL = gastrocnemius lateralis, SO = soleus.

**Figure 3.**
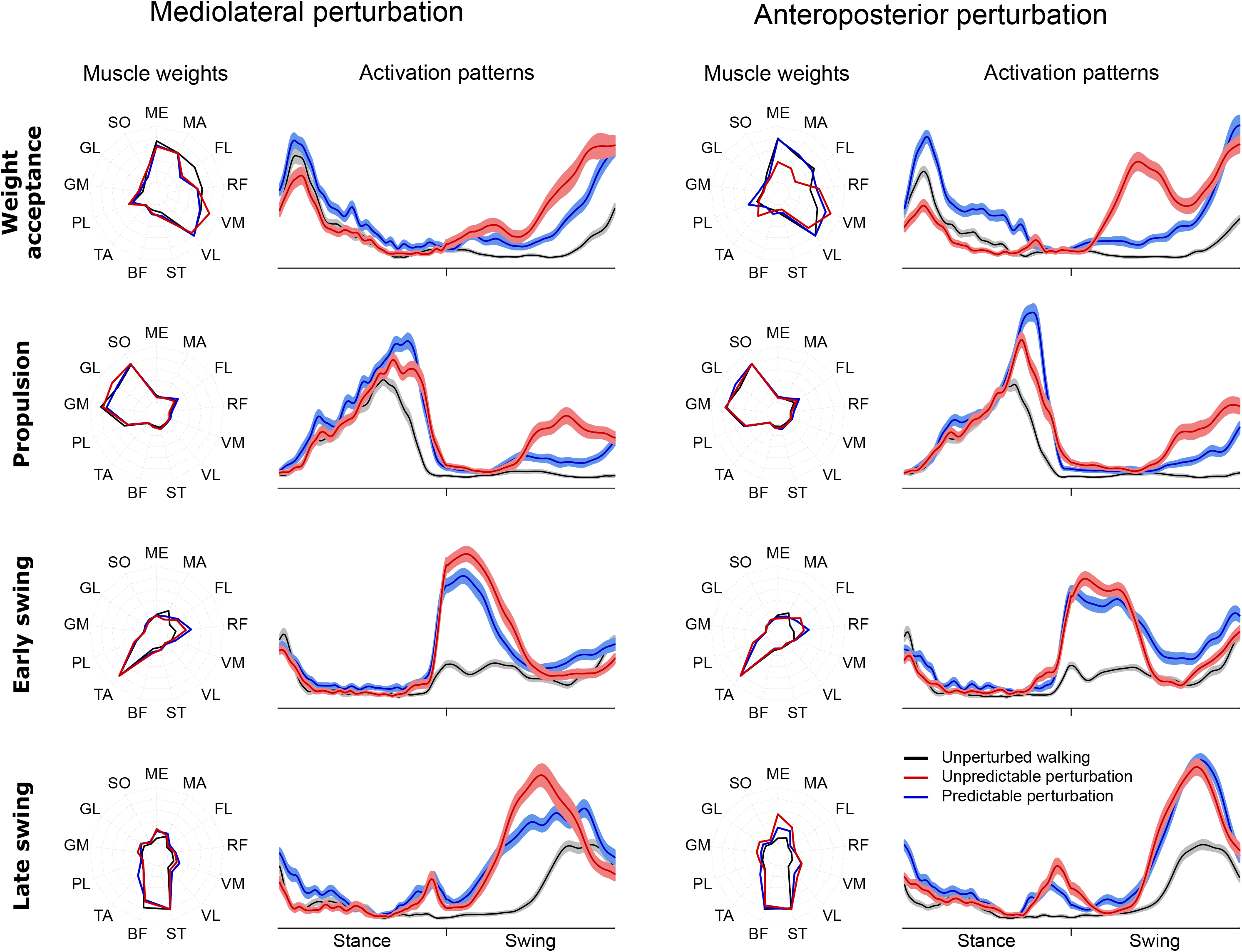
Means of the muscle weights and means ± standard error of the activation patterns of the four fundamental muscle synergies during unperturbed walking (black line), unpredictable (red line) and predictable (blue line) perturbations. The amplitudes of the activation patterns and muscle weights were normalized to 1. The stance and swing phases were normalized to an equal distribution of data points. ME = gluteus medius, MA = gluteus maximus, FL = tensor fasciæ latæ, RF = rectus femoris, VM = vastus medialis, VL = vastus lateralis, ST = semitendinosus, BF = biceps femoris, TA = tibialis anterior, PL = peroneus longus, GM = gastrocnemius medialis, GL = gastrocnemius lateralis, SO = soleus.

We found a statistically significant effect of gait cycle (p < 0.007) on the CS of the activation patterns (figure 4) and muscle weights (figure 5) in all synergies, which shows a modulation of the spatiotemporal structure of the muscle synergies during and after the perturbation (figures 6). The CS of the activation patterns decreased (p < 0.001) in all synergies and conditions during the perturbation cycle compared to the unperturbed one (figure 4). The CS of the muscle weights was also significantly decreased (p < 0.04) in most of the synergies during the perturbation and the first recovery cycle (figure 5). Furthermore, there was a statistically significant effect of predictability (p < 0.05) on almost all CS of the activation patterns and muscle weights with lower values in the unpredictable condition in both the mediolateral and anteroposterior direction (figures 4 and 5). Only in the mediolateral weight acceptance synergy of the muscle weights, the decrease of CS in the predictable condition was not statistically significant (p = 0.12), while a significant (p < 0.001) interaction effect of predictability by gait cycle was found in that synergy. The significant effect of predictability on the similarity of the activation patterns was present until the third-fourth recovery cycle (figure 4).

**Figure 4.**
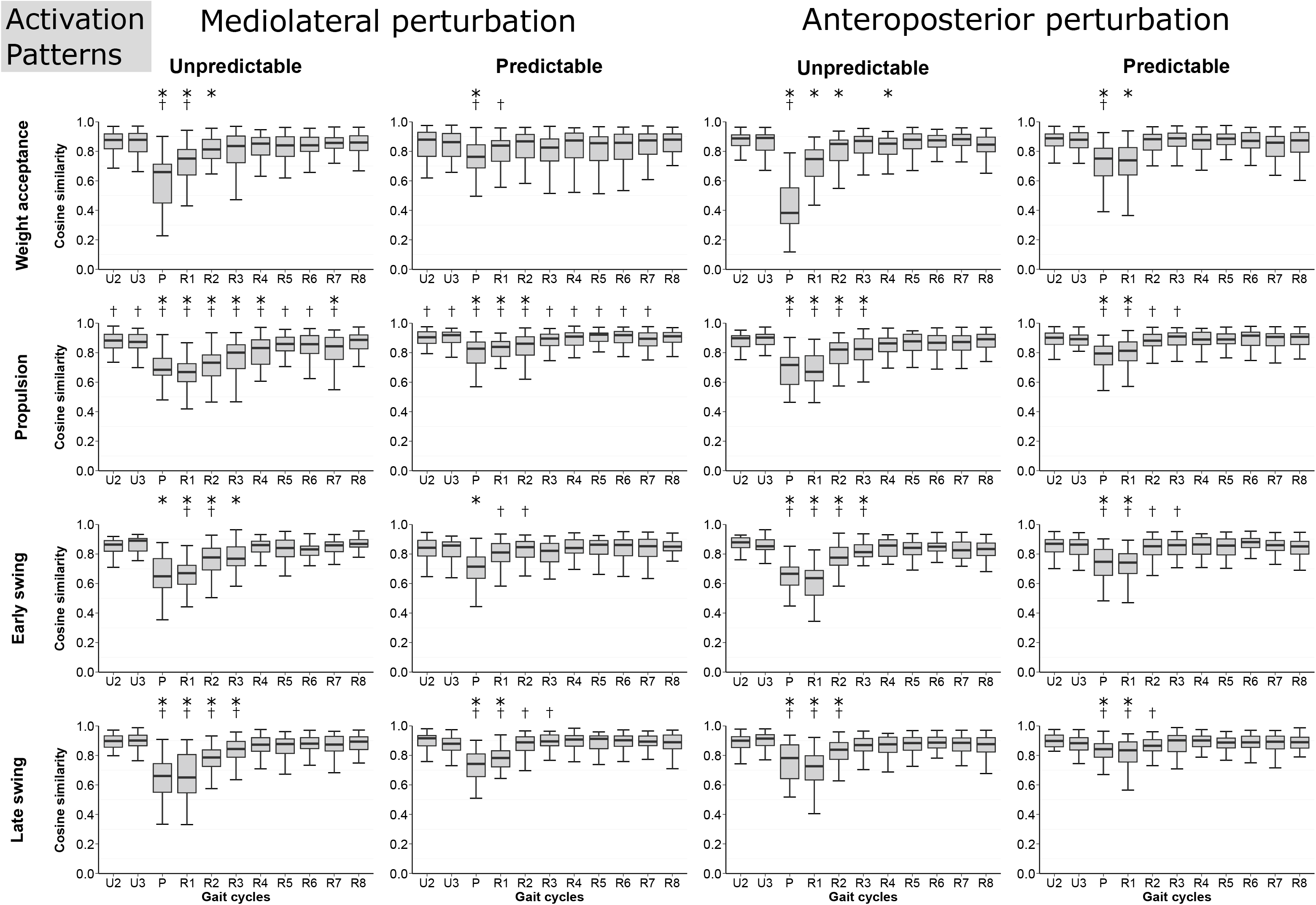
Cosine similarity (CS) of the activation patterns of the muscle synergies for the two unperturbed cycles (U2, U3), the perturbation cycle (P) and the following 8 recovery cycles (R1 to R8) with regard to the first unperturbed cycle. Statistically significant differences to the CS of the unperturbed cycle (U2) (*) and between the unpredictable and predictable condition (†) are highlighted (p<0.05).

**Figure 5.**
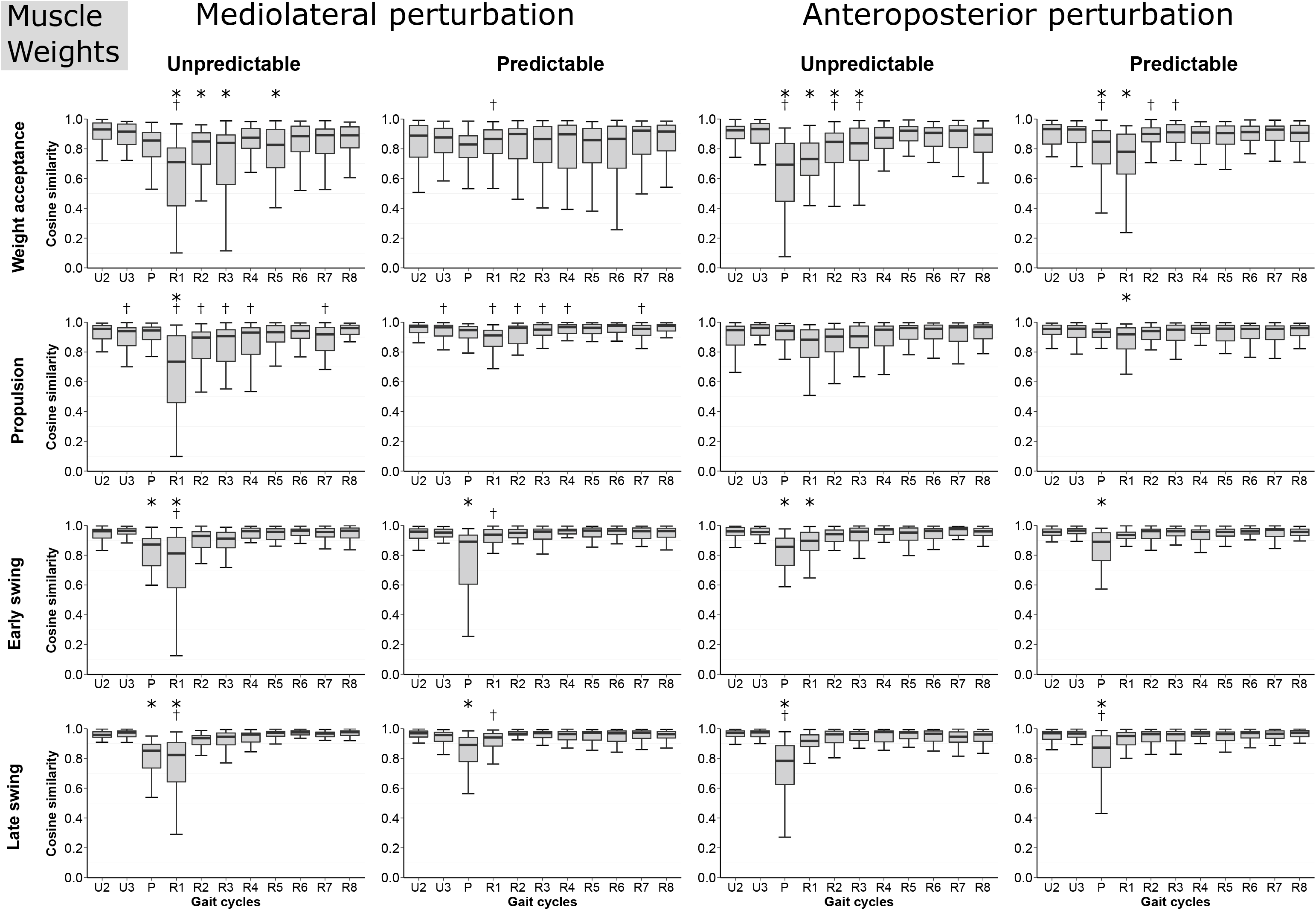
Cosine similarity (CS) of the muscle weights of the muscle synergies for the two unperturbed cycles (U2, U3), the perturbation cycle (P) and the following 8 recovery cycles (R1 to R8) with regard to the first unperturbed cycle. Statistically significant differences to the CS of the unperturbed cycle (U2) (*) and between unpredictable and predictable condition (†) are highlighted (p<0.05).

**Figure 6.**
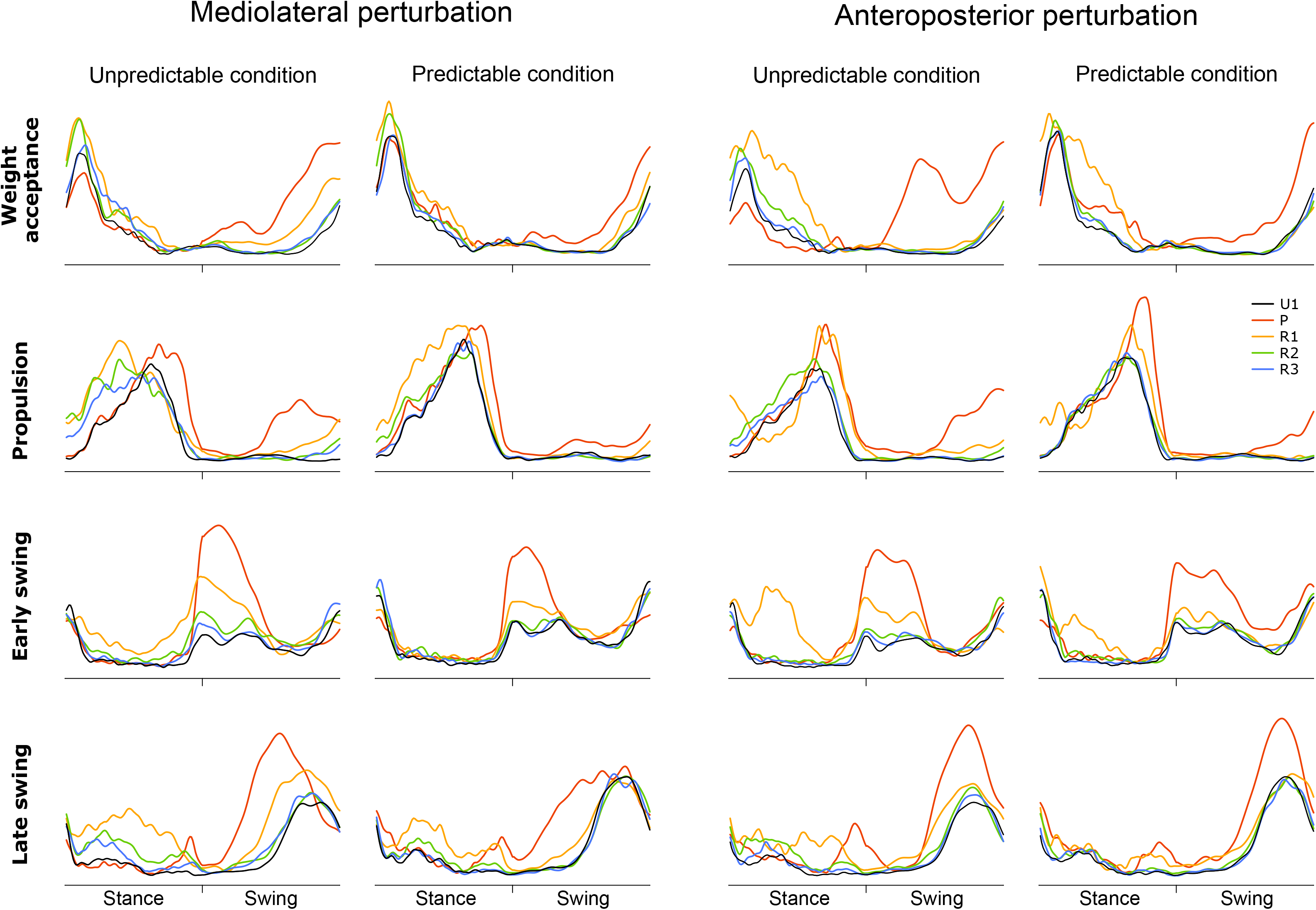
Mean curves of the activation patterns of the muscle synergies for the four different conditions during the unperturbed walking (U1), the perturbation cycle (P) and the following three recovery cycles (R1 to R3). All amplitudes of the activation patterns were normalized to 1. The stance and swing phases were normalized to an equal distribution of data points.

We found a significant effect of cycle (p < 0.02) and predictability (p < 0.02) on the CoA for nearly all synergies in both mediolateral and anteroposterior direction (table 2). Only the early swing synergy showed no significant effect of predictability (p < 0.349), but an interaction effect of cycle by predictability (p < 0.001). In the perturbation and the first recovery cycle, the CoA of the activation patterns shifted either earlier or later within the gait cycle compared to the unperturbed walking with greater changes in the unpredictable condition (table 2). The shift of the CoA was statistically significant in all synergies and conditions (p < 0.05) except for the late swing synergy in the predictable anteroposterior condition. The alterations in the CoA lasted until the 5th recovery cycle (table 2). There was an effect of gait cycle (p < 0.005) and a cycle by predictability interaction (p < 0.005) on the FWHM of nearly all synergies in both the mediolateral and anteroposterior direction. In most cases, the FWHM of the activation patterns became wider (table 3). The significant differences compared to the unperturbed walking were mainly observed in the perturbation and in the first recovery cycle for all investigated conditions. The increased widening of the activation patterns during the perturbation and first recovery cycle was in most cases larger in the unpredictable condition (table 3). There was a decrease in FWHM of the propulsion synergy in the first recovery cycle after the unpredictable anteroposterior condition (table 3).

**Table 2.** Means ± SD values of the center of activity (CoA) of the activation patterns of the muscle synergies during the three unperturbed cycles (U1 to U3), the perturbation cycle (P) and the following 8 recovery cycles (R1 to R8) for the four different conditions. Statistically significant differences to the first unperturbed cycle (*) and between the unpredictable and predictable condition (†) are highlighted (p<0.05).

| COA [time points] |  |  | U1 | U2 | U3 | P | R1 | R2 | R3 | R4 | R5 | R6 | R7 | R8 |
| --- | --- | --- | --- | --- | --- | --- | --- | --- | --- | --- | --- | --- | --- | --- |
| Mediolateral perturbation | Unpredictable | Weight acceptance | 20.6 ± 26.8 | 20.2 ± 27.7 | 26.6 ± 43.0 | 152.1 ± 69.7** | 44.1 ± 67.7** | 25.0 ± 39.8 | 17.1 ± 9.2 | 17.1 ± 7.4 | 20.8 ± 26.7 | 19.8 ± 26.8 | 27.3 ± 43.2 | 17.6 ± 8.4 |
| | | Propulsion | 54.9 ± 7.0 | 54.7 ± 5.7 | 53.7 ± 6.4 | 56.5 ± 24.6 | 38.9 ± 9.3** | 41.0 ± 9.5** | 46.1 ± 8.4** | 50.0 ± 6.9** | 53.1 ± 6.3 | 53.3 ± 8.3 | 53.5 ± 7.9 | 54.1 ± 6.5 |
| | | Early swing | 162.4 ± 19.1 | 160.0 ± 19.6† | 159.2 ± 13.9 | 120.9 ± 19.4* | 114.1 ± 33.8** | 136.1 ± 37.5** | 152.2 ± 32.6 | 155.1 ± 33.2 | 158.4 ± 20.1 | 150.7 ± 32.8 | 153.1 ± 17.9 | 154.0 ± 37.6 |
| | | Late swing | 166.9 ± 50.1 | 158.3 ± 59.2 | 163.7 ± 54.2 | 156.6 ± 8.7 | 112.3 ± 83.9* | 124.9 ± 82.4** | 164.2 ± 54.3 | 157.9 ± 60.8 | 152.7 ± 68.0 | 170.7 ± 44.4 | 160.9 ± 58.1 | 165.4 ± 53.1 |
| | Predictable | Weight acceptance | 17.2 ± 9.7 | 18.4 ± 25.1 | 16.1 ± 8.2 | 70.7 ± 84.0** | 17.1 ± 8.3† | 17.3 ± 8.2 | 24.2 ± 27.4 | 17.0 ± 7.3 | 18.9 ± 7.8 | 16.4 ± 7.1 | 15.9 ± 7.1 | 17.3 ± 6.8 |
| | | Propulsion | 55.3 ± 7.4 | 55.3 ± 6.5 | 54.0 ± 5.9 | 57.1 ± 11.2 | 48.4 ± 6.5** | 50.5 ± 7.1** | 55.6 ± 8.7† | 55.6 ± 6.9† | 55.3 ± 6.1 | 56.6 ± 6.8 | 57.2 ± 6.8 | 56.8 ± 5.0 |
| | | Early swing | 158.1 ± 27.9 | 147.3 ± 42.2**† | 153.8 ± 28.4 | 126.8 ± 19.8* | 148.5 ± 29.9† | 153.2 ± 32.4† | 160.6 ± 29.7 | 152.1 ± 40.2 | 152.0 ± 33.8 | 156.4 ± 29.9 | 150.2 ± 36.7 | 157.8 ± 27.6 |
| | | Late swing | 169.5 ± 45.8 | 169.1 ± 44.9 | 165.1 ± 52.4 | 164.9 ± 12.9 | 131.3 ± 79.0* | 158.0 ± 65.4† | 169.1 ± 48.7 | 172.8 ± 42.1 | 168.5 ± 48.5 | 170.5 ± 46.1 | 170.2 ± 46.3 | 164.6 ± 53.7 |
| Anteroposterior perturbation | Unpredictable | Weight acceptance | 20.6 ± 25.7 | 31.9 ± 53.9† | 22.3 ± 34.2 | 155.1 ± 43.2** | 24.2 ± 8.6 | 17.3 ± 8.1 | 29.2 ± 46.9 | 27.6 ± 41.7 | 25.9 ± 44.4 | 17.0 ± 8.0 | 18.5 ± 26.5 | 16.6 ± 9.4 |
| | | Propulsion | 54.0 ± 6.1 | 52.7 ± 7.2 | 54.4 ± 5.5 | 49.7 ± 28.2** | 56.8 ± 9.5 | 49.3 ± 6.4** | 50.9 ± 8.4 | 51.8 ± 7.2 | 51.0 ± 8.2 | 52.9 ± 8.4 | 53.6 ± 7.2 | 53.2 ± 5.8 |
| | | Early swing | 163.9 ± 17.2 | 158.5 ± 13.8 | 154.5 ± 30.7 | 124.6 ± 16.7** | 82.3 ± 60.2** | 158.8 ± 29.1 | 156.8 ± 18.2 | 158.6 ± 16.9 | 159.2 ± 20.2 | 162.9 ± 14.1 | 160.3 ± 18.9 | 158.2 ± 29.2 |
| | | Late swing | 176.5 ± 33.5 | 167.8 ± 49.9 | 172.9 ± 41.2 | 168.9 ± 25.1 | 138.0 ± 74.1**† | 143.7 ± 75.9* | 154.2 ± 64.7 | 146.4 ± 70.5* | 155.5 ± 65.5 | 159.5 ± 60.3 | 161.0 ± 58.0 | 163.5 ± 57.3 |
| | Predictable | Weight acceptance | 15.0 ± 7.1 | 16.9 ± 7.5† | 16.7 ± 7.7 | 55.5 ± 76.3** | 24.4 ± 8.7 | 16.8 ± 6.1 | 17.0 ± 6.8 | 18.1 ± 7.3 | 16.7 ± 7.2 | 19.0 ± 7.6 | 19.0 ± 6.7 | 18.1 ± 7.4 |
| | | Propulsion | 54.2 ± 5.7 | 54.0 ± 4.9 | 53.9 ± 6.4 | 60.1 ± 14.4** | 55.3 ± 9.7 | 54.1 ± 7.1† | 53.7 ± 6.0 | 54.1 ± 6.4 | 54.5 ± 5.8 | 55.3 ± 5.6 | 54.8 ± 6.4 | 56.1 ± 6.5 |
| | | Early swing | 157.3 ± 20.2 | 155.9 ± 26.3 | 151.4 ± 25.7 | 140.2 ± 15.3** | 117.3 ± 65.7**† | 147.8 ± 43.9 | 154.8 ± 18.1 | 161.1 ± 19.6 | 150.3 ± 42.4 | 159.1 ± 17.6 | 155.8 ± 18.7 | 160.7 ± 15.9 |
| | | Late swing | 180.2 ± 24.4 | 172.3 ± 42.0 | 170.4 ± 46.2 | 174.5 ± 24.6 | 168.5 ± 47.4† | 164.4 ± 56.3 | 162.2 ± 58.3 | 166.9 ± 52.2 | 160.7 ± 59.0 | 168.6 ± 51.7 | 169.2 ± 50.4 | 167.8 ± 51.4 |

**Table 3.** Means ± SD values of the full width at half maximum (FWHM) of the activation patterns of the muscle synergies during the three unperturbed cycles (U1 to U3), the perturbation cycle (P) and the following 8 recovery cycles (R1 to R8) for the four different conditions. Statistically significant differences to the first unperturbed cycle (*) between the unpredictable and predictable condition (†) are highlighted (p<0.05).

| FWHM [time points] |  |  | U1 | U2 | U3 | P | R1 | R2 | R3 | R4 | R5 | R6 | R7 | R8 |
| --- | --- | --- | --- | --- | --- | --- | --- | --- | --- | --- | --- | --- | --- | --- |
| Mediolateral perturbation | Unpredictable | Weight acceptance | 20.5 ± 8.2 | 21.2 ± 8.4 | 18.6 ± 8.2 | 33.5 ± 15.9* | 26.9 ± 12.2* | 17.1 ± 7.7 | 22.8 ± 10.5 | 17.9 ± 6.6 | 19.0 ± 8.6 | 20.8 ± 10.4 | 22.2 ± 9.9 | 18.9 ± 8.8 |
| | | Propulsion | 30.4 ± 10.8 | 28.2 ± 11.2 | 32.4 ± 11.6 | 45.4 ± 20.1** | 30.1 ± 10.9<sup>†</sup> | 34.4 ± 13.6 | 35.2 ± 12.2 | 33.2 ± 14.2 | 31.6 ± 12.4 | 31.6 ± 10.8 | 29.0 ± 11.2 | 30.7 ± 11.8 |
| | | Early swing | 36.6 ± 22.4 | 32.1 ± 17.0 | 34.9 ± 17.8 | 37.2 ± 13.3 | 39.3 ± 17.3 | 40.5 ± 17.5 | 35.5 ± 17.6 | 37.1 ± 22.5 | 33.6 ± 16.4 | 33.8 ± 16.4 | 34.0 ± 18.7 | 33.2 ± 17.4 |
| | | Late swing | 33.8 ± 13.0 | 32.6 ± 12.0 | 30.7 ± 10.8 | 38.9 ± 16.1<sup>†</sup> | 49.0 ± 19.4** | 34.8 ± 13.4 | 38.2 ± 15.2 | 35.0 ± 15.2 | 32.8 ± 12.9 | 33.3 ± 11.6 | 32.3 ± 11.2 | 33.4 ± 10.2 |
| | Predictable | Weight acceptance | 22.5 ± 10.3 | 20.2 ± 9.2 | 20.1 ± 9.4 | 33.7 ± 15.5* | 22.2 ± 12.1 | 21.5 ± 8.4 | 20.3 ± 8.7 | 21.4 ± 9.2 | 21.3 ± 10.0 | 18.2 ± 7.8 | 18.8 ± 7.4 | 20.2 ± 9.0 |
| | | Propulsion | 31.3 ± 10.8 | 29.4 ± 10.6 | 31.1 ± 11.8 | 37.4 ± 18.2** | 38.7 ± 14.6** | 36.4 ± 13.4 | 30.1 ± 13.1 | 28.9 ± 12.6 | 29.8 ± 11.1 | 31.0 ± 13.2 | 30.8 ± 11.6 | 30.9 ± 10.6 |
| | | Early swing | 34.1 ± 18.2 | 39.4 ± 23.5 | 39.2 ± 20.0 | 36.5 ± 17.6 | 44.2 ± 19.1 | 34.0 ± 21.4 | 33.7 ± 20.1 | 38.2 ± 20.8 | 35.6 ± 17.8 | 37.5 ± 22.3 | 36.1 ± 21.5 | 36.6 ± 19.5 |
| | | Late swing | 33.4 ± 11.5 | 33.8 ± 12.2 | 31.2 ± 13.1 | 47.5 ± 18.2** | 37.3 ± 15.0<sup>†</sup> | 36.4 ± 14.2 | 33.6 ± 9.9 | 31.6 ± 8.9 | 33.1 ± 11.2 | 32.5 ± 10.2 | 32.2 ± 9.4 | 32.1 ± 9.7 |
| Anteroposterior perturbation | Unpredictable | Weight acceptance | 20.1 ± 12.5 | 21.5 ± 10.4 | 20.8 ± 9.4 | 38.7 ± 16.2** | 24.0 ± 10.6 | 21.2 ± 10.0 | 21.4 ± 7.2 | 19.0 ± 8.7 | 19.4 ± 9.6 | 19.8 ± 8.8 | 19.2 ± 9.8 | 21.4 ± 10.2 |
| | | Propulsion | 31.1 ± 13.2 | 31.6 ± 13.8 | 29.9 ± 10.0 | 35.1 ± 18.9<sup>†</sup> | 22.5 ± 7.3** | 32.8 ± 12.1 | 32.4 ± 13.3 | 32.8 ± 13.2 | 33.9 ± 12.6 | 31.0 ± 12.6 | 30.9 ± 11.2 | 31.4 ± 11.4 |
| | | Early swing | 31.9 ± 21.4 | 37.7 ± 20.9 | 30.8 ± 16.2 | 39.2 ± 14.8<sup>†</sup> | 32.3 ± 16.0 | 31.9 ± 21.2 | 35.3 ± 19.1 | 35.5 ± 19.7 | 35.7 ± 20.0 | 32.3 ± 23.7 | 33.7 ± 21.6 | 35.8 ± 25.2 |
| | | Late swing | 32.5 ± 13.1 | 33.1 ± 13.0 | 31.7 ± 10.6 | 36.5 ± 12.8 | 40.6 ± 15.9* | 37.1 ± 15.1 | 33.0 ± 11.5 | 36.1 ± 13.3 | 32.3 ± 12.1 | 32.9 ± 10.6 | 32.1 ± 10.5 | 31.8 ± 12.7 |
| | Predictable | Weight acceptance | 19.7 ± 7.4 | 19.6 ± 6.7 | 19.5 ± 10.3 | 29.8 ± 11.7** | 25.1 ± 9.3* | 20.5 ± 9.0 | 22.3 ± 11.5 | 20.2 ± 10.1 | 18.4 ± 8.1 | 21.0 ± 9.6 | 20.5 ± 9.1 | 19.6 ± 7.7 |
| | | Propulsion | 33.0 ± 12.2 | 30.5 ± 9.7 | 32.9 ± 11.2 | 27.9 ± 15.3<sup>†</sup> | 29.6 ± 11.5<sup>†</sup> | 32.4 ± 12.2 | 32.3 ± 13.3 | 32.1 ± 11.6 | 31.9 ± 12.8 | 31.8 ± 10.8 | 31.9 ± 11.1 | 31.4 ± 10.1 |
| | | Early swing | 30.9 ± 18.0 | 37.0 ± 21.1 | 37.6 ± 18.0 | 48.1 ± 19.8** | 28.8 ± 19.7 | 40.5 ± 17.1 | 35.9 ± 17.9 | 34.5 ± 20.0 | 32.5 ± 17.0 | 32.2 ± 21.2 | 37.4 ± 20.7 | 32.2 ± 18.8 |
| | | Late swing | 31.7 ± 11.9 | 34.0 ± 11.5 | 33.6 ± 12.0 | 39.2 ± 15.8* | 37.7 ± 16.3 | 33.3 ± 12.0 | 31.3 ± 10.9 | 32.2 ± 9.9 | 34.0 ± 11.0 | 32.3 ± 13.0 | 33.2 ± 10.8 | 30.7 ± 9.7 |

The SPM analysis showed significant differences (p < 0.05) of EMG-activity before the initiation of the perturbation between the predictable and unpredictable conditions (figure 7). In both mediolateral and anteroposterior perturbations, rectus femoris, vastus medialis and vastus lateralis showed a significantly higher EMG-activity (p ≤ 0.002) before the perturbation in the predictable compared to the unpredictable condition. During the mediolateral perturbations, biceps femoris (p = 0.049), peroneus longus (p < 0.001) and gastrocnemius lateralis (p < 0.001) showed a significantly higher EMG-activity in the predictable condition, whereas in the anteroposterior perturbations, the EMG-activity of semitendinosus (p = 0.006), tibialis anterior (p = 0.003) and gastrocnemius medialis (p = 0.033) was higher (figure 7).

**Figure 7.**
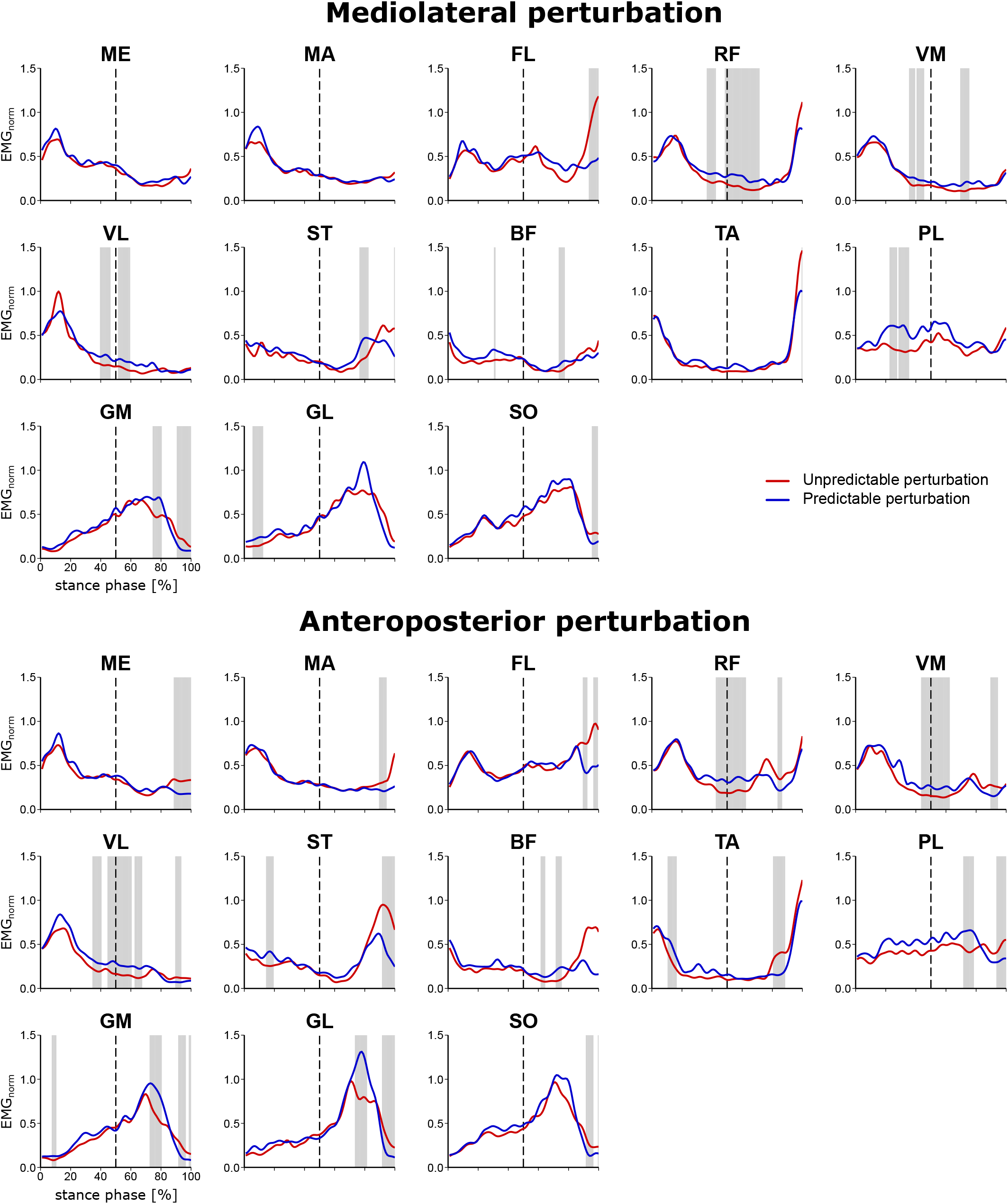
Mean curves of the electromyographic (EMG) activity of the 13 recorded leg muscles during the stance phase of the perturbed gait cycle for the unpredictable (red line) and predictable condition (blue line). The start of the perturbation is indicated by a dashed line. For each muscle the amplitude was normalized to the maximum EMG activity during unperturbed walking. The grey vertical bands highlight ranges of significant differences (p<0.05) between the curves based on statistical parametric mapping. ME = gluteus medius, MA = gluteus maximus, FL = tensor fasciæ latæ, RF = rectus femoris, VM = vastus medialis, VL = vastus lateralis, ST = semitendinosus, BF = biceps femoris, TA = tibialis anterior, PL = peroneus longus, GM = gastrocnemius medialis, GL = gastrocnemius lateralis, SO = soleus.

## Discussion

In the current study, we investigated the effects of predictable and unpredictable perturbations on the modular organization of locomotion in older adults. For both the muscle weights and activation patterns, we found a significant decrease in CS between the perturbed cycle and the following recovery cycles compared to unperturbed walking, indicating a modulation of the spatiotemporal structure of muscle synergies by the sensorimotor system in order to maintain gait stability after the perturbations. The number of muscle synergies remained unchanged in both predictable and unpredictable perturbations and the four basic walking synergies (weight acceptance, propulsion, early swing and late swing) were preserved, confirming our first hypothesis. Further, we found a smaller decrease in CS during the predictable compared to the unpredictable perturbations, indicating a less pronounced modulation of the spatiotemporal structure of muscle synergies, confirming our second hypothesis. The findings were generally consistent for both anteroposterior and mediolateral perturbations and thus invariant to the type of perturbation.

The reduced CS in all activation patterns in the perturbed gait cycle evidenced a clear modification of the temporal components of the muscle synergies. Similar to the activation patterns, there was a modification of the muscle weights during the perturbed cycle in most of the cases, indicating a flexible organization also of the spatial components of muscle synergies during perturbed gait. Although the spatiotemporal structure of muscle synergies was modified during gait perturbations, the basic structure and composition was preserved and the four basic synergies found in unperturbed walking were enough to counteract the challenging perturbations in both predictable and unpredictable trials. The fixed set of muscle synergies was maintained also during the mediolateral perturbations, whereas the gait biomechanics deviated from the unperturbed fore-aft walking. During the mediolateral gait perturbations, a fast lateral step or a crossing step is needed to counteract a sideways fall (Madehkhaksar et al., 2018). Our results showed that the participants’ acute lateral steps were performed by modulating the basic activation patterns of the four synergies for unperturbed walking, leaving the number of synergies unchanged. These findings are in line with previous studies investigating continuous and less intense perturbations during walking (Martino et al., 2015; Santuz et al., 2018a), where preservation of the number of synergies was observed.

The recruitment of the four basic walking synergies in the perturbed and recovery gait cycles indicates a robust synergy structure and a simplification of postural stability control by using activation patterns of a few and well-known muscle synergies with specific adjustments within the synergies. Stability control via recruitment of existing muscle synergies during perturbed walking with the integration of somatosensory information may reduce the need for higher-level control and promote the transfer of adaptation between different kinds of perturbations (König et al., 2022). A recent study in rats reported a robust spinal structure of muscle synergies during walking, which was preserved despite different developmental and learning processes (Yang et al., 2019). Takei et al. (2017) showed that spinal premotor interneurons generate the spatiotemporal components of muscle synergies during a precision motor task in monkeys and suggested that superimposed corticomotoneuronal pathways are responsible for more fractionated motor control. The gait perturbations applied in the current study increased the risk of falling and triggered fast reactive recovery responses, which might require less involvement of the corticomotoneuronal system. The recruitment of shared synergies for the control of different motor tasks has been reported in the past (D’Avella & Bizzi, 2005) and was suggested to be a general strategy of the neuromotor system to flexibly and efficiently create complex motor behavior. Using a simulation approach, d’Avella and Pai (2010) found that a modular controller was able to adapt faster to applied force perturbations if the neuromotor control system used a set of pre-existing muscle synergies. Berger et al. (2013) reported higher adaptation rates in a novel motor task that can be executed with pre-existing synergies compared to a task that required new muscle synergies. Chvatal and Ting (2012) found modifications in the temporal recruitment of common locomotor muscle synergies during experienced gait perturbations in young participants. Our study expands those findings showing that older adults used a fixed number of muscle synergies and adjusted their temporal recruitments to maintain stability in both predictable and unpredictable gait perturbations with and without prior experience. The activation of pre-existing muscle synergies and adjustment of the time of their recruitment may therefore provide an effective way to counteract locomotor perturbations, where fast reactive responses are necessary to maintain postural stability (Bohm et al., 2015; Karamanidis et al., 2020; Zhang et al., 2020).

The recruitment of muscle synergies in the perturbed gait cycles was associated with specific functional phases of walking and their modulation corresponded to the perturbation-specific stability requirements. In our experimental design, the perturbations were introduced in the middle of the stance phase with a quick shift of the treadmill’s belt in anteroposterior or mediolateral direction. The activation patterns of the early swing synergy showed a steep increase with a higher contribution of the tibialis anterior and rectus femoris in the perturbed cycles. Further, the shift of the CoA to an earlier time point following the perturbation in this synergy suggests a faster recruitment of the early swing synergy as a reactive control mechanism in order to increase the efficacy of execution of the following step (König et al., 2022). The first step after a gait perturbation is crucial for balance recovery and fall prevention, especially in older individuals where reactive responses are deteriorated (Arampatzis et al., 2008; Bierbaum et al., 2011; Mersmann et al., 2013). A fast and long step after the perturbation increases the base of support and results in an improvement of stability (Bierbaum et al., 2010, 2011; Maki & McIlroy, 2006). Similar to the early swing synergy, the activation pattern of the propulsion synergy showed a rapid increase in the second phase of stance in the perturbed cycles. Taken into consideration that in this synergy the plantar flexor muscles are the main contributors and achieved high values in their muscle weights compared to other muscles, the steep increase of the activation pattern in the propulsion synergy indicates a rapid force development of the plantar flexor muscles after the initiation of the perturbation, which is functionally relevant to prevent a fall (Pijnappels et al., 2005; Robinovitch et al., 2002). The rapid increase in the basic activation patterns of the propulsion synergy is functionally associated with the earlier plantar flexion after anteroposterior and mediolateral gait perturbations compared to unperturbed walking (Taborri et al., 2022). Further, the widening of the propulsion synergy corresponds to a longer activation and thus longer force generation of the plantar flexor muscles relative to the stance phase during propulsion, indicating a mechanism for an effective push-off and initiation of the swing phase. It is to mention that muscle force generation is also dependent on muscle contractile conditions like muscle force potentials due to the force-length and force-velocity relationships. Nevertheless, a widening of the activation of the triceps surae muscles in spite of varying force-length-velocity potentials will introduce a longer force generation and thus a more effective push-off.

The widening of the basic activation patterns of the muscle synergies during the perturbed and the first recovery steps lead to an increased overlap of activation between chronologically adjacent synergies and can be interpreted as a compensatory mechanism adopted by the central nervous system to deal with the perturbation and promote robust postural control (Janshen et al., 2020; Santuz et al., 2018a). Perturbations clearly affected gait stability, as shown by the modified temporal gait parameters (i.e. cadence, stance and swing times). Cadence was increased and stance and swing times were decreased in the perturbed and in the following recovery steps, demonstrating biomechanical adjustments to regain stability after the perturbation. Maintenance of stability is a primary task-relevant variable during locomotion (Cajigas et al., 2017) and, therefore, corrective locomotor actions are important after perturbations (Bohm et al., 2015; Iturralde & Torres-Oviedo, 2019). Widening of the basic activation patterns of the muscle synergies following perturbations as a strategy to provide robust motor control seems to be a common mechanism in biological systems. Widening of activation patterns during challenging locomotor tasks has been found in healthy individuals (Martino et al., 2015; Santuz et al., 2018a), multiple sclerosis patients (Janshen et al., 2020, 2021) and in mice (Santuz et al., 2022).

The observed modulations in the spatiotemporal structure of muscle synergies during the perturbed cycle were less pronounced in the predictable condition. Prior experience with specific perturbations can promote proactive adjustments of motor control (Bohm et al., 2015; Patla, 2003) and reduce the consequences of a perturbation (Bierbaum et al., 2010; Kanekar & Aruin, 2015; Marigold & Patla, 2002). The sensorimotor system can prepare a specific motor action in a predictive manner before its execution by building up an expectation of sensory inputs and then comparing the expected and actual inputs in order to successfully drive the desired movement (Wolpert, 1997). Prior experience with a perturbation can also improve the corrective responses after the perturbation (Bierbaum et al., 2011; Van Der Linden et al., 2007) possible through a supraspinal mediation of sensory inputs. We found an increased EMG-activity in several lower leg muscles before the initiation of the perturbation (i.e. middle of the stance phase) in the predictable compared to the unpredictable condition, which indicates proactive adjustments in the activation patterns. During the predictable condition, all knee extensors showed increased activity within the last ∼100 ms prior to the perturbation in both anteroposterior and mediolateral directions. This might have contributed to an improved limb support of the stance leg increasing hip height (Pai et al., 2010), as an important strategy of the neuromotor system to avoid falling (Pavol & Pai, 2007). An increased limb support could further benefit a fast initiation of the recovery step to improve postural stability (Bierbaum et al., 2010; Maki & McIlroy, 2006).

In conclusion, our findings demonstrated that older adults used the same basic structure and composition of the four walking muscle synergies, yet modulated their spatiotemporal components in order to counteract gait perturbations. The above behavior was consistent during the perturbed and the recovery steps and in all applied walking perturbations (i.e. anteroposterior, mediolateral, unpredictable, predictable). The selection of pre-existing muscle synergies and adjustment of their recruiting timing indicates a flexible postural control in human gait during challenging locomotor conditions, which may facilitate the effectiveness in coping with perturbations and promote the transfer of adaptation between different types of perturbations.

## Acknowledgments

We like to thank all participants for their commitment to this study. We further thank Carlotta Körbi, Jaqueline Martini and Thomas Gerhardy for their assistance in data collection and Arno Schroll for statistical consultation.

## Author contributions

Conceptualization: L.B., A.A.; Methodology: L.B., A.S., A.A.; Software: L.B., A.S.; Validation: L.B., A.S.; Formal analysis: L.B., A.S., A.A.; Investigation: L.B.; Resources: M.S., A.A.; Data Curation: L.B., A.S.; Writing – original draft preparation: L.B., A.A.; Writing – review and editing: L.B., A.S., F.M., S.B., M.S., A.A.; Visualization: L.B.; Supervision: A.A.; Project administration: L.B., M.S., A.A.; Funding acquisition: M.S., A.A.

